# Land-use preferences of the European green toad (*Bufotes viridis*) in the city of Vienna (Austria): the importance of open land in urban environments

**DOI:** 10.1101/2022.07.05.498873

**Authors:** Lukas Landler, Stephan Burgstaller, Silke Schweiger

## Abstract

Urban areas are increasing worldwide, which poses treats to animal wildlife. However, in certain cases cities can provide refuges for endangered animals. The European green toad (*Bufotes viridis*) is one of such examples, which is known from cities throughout their distribution. In contrast, considerable areas of their former (primary) habitats have been degraded. The primary habitats of this species include steppes and wild river floodplains, both characterized by dynamic changes and the presence of open areas. We used available green toad observation data (2007-2020) to model the effects of land-use types on occurrence probability in the city of Vienna. Forest and densely populated areas were highly significantly negatively associated with green toad presence, while transformation/construction site areas showed a strong positive effect. Such occurrence pattern might be characteristic for early succession species, which depend on stochastic environmental disturbances (e.g., droughts and floods) in their primary habitats. We argue that urban landscape planning should appreciate the potential ecological value of open land in cities which is either in a transition phase or a permanent wasteland. Ecological managing of such landscape could vastly increase urban biodiversity.

## Background

Urban areas as well as human population densities in cities are increasing worldwide (Angel et al., 2011; Gao & O’Neill, 2020). Such developments are threatening plant and animal wildlife and can also pose risks for humans living in cities (e.g., wildlife related diseases, animals as disease vectors and direct interactions with animals) (Bradley & Altizer, 2007; Hassell et al., 2017; Luck & Smallbone, 2010; McKinney, 2008; Moore et al., 2003; Murray et al., 2019; Vlahov, 2002). Often urban areas are dominated by relatively few urban exploiter species, which thrive under such conditions, taking advantage of food supply and reduced species competition or predation risk (i.e., rock pigeon, brown rat, house sparrow) (Marzluff, 2001; McKinney, 2002). On the other hand, while in rural areas agricultural practices are intensifying and road networks are expanding, urban green spaces can provide refuge for threatened and otherwise rare animals (D. M. Hall et al., 2017; Soanes & Lentini, 2019). Van Helden et al. (2021) found that, in South-West Australia western ringtail possums (*Pseudocheirus occidentalis*) can survive exclusively in residential gardens with no noticeable detrimental effects. In an expanding urban area in Ghana Ofori et al. (2018) showed that urban green areas can conserve large proportions of the native small mammal biodiversity.

However, while some animal taxa, such as birds and small mammals, have several urban exploiters in their midst, other species struggle considerably with human transformations. Amphibians, for example, are under threat globally. It is estimated that currently over 40 % of amphibian species are threatened or endangered (IUCN, 2021). Factors contributing to amphibian decline include fungal disease, pollution, land degradation as well as urbanization (Beebee & Griffiths, 2005; Hamer & McDonnell, 2008; Scheffers & Paszkowski, 2011). For many amphibian species, increased drainage and the resulting loss of natural water bodies leads to rapid population declines (Collins & Storfer, 2003; Rannap et al., 2007; Suislepp et al., 2011). Amphibian species richness usually declines with increasing urbanization because anthropogenically altered environments often fail to provide suitable breeding sites as well as terrestrial habitats (Hamer & McDonnell, 2008). However, urban parks for example can provide an oasis for specialized amphibian species and contribute to amphibian conservation (Brand & Snodgrass, 2010; Scheffers & Paszkowski, 2011). For instance, the Sydney Olympic Park harbors the largest known population of the endangered green and golden bell frog (*Litoria aurea*) (Darcovich & O’Meara, 2008). The European green toad (*Bufotes viridis*) is another amphibian species, that is well-known for the presence in urban areas (Bogdan, 2014; Konowalik et al., 2020; Kühnel & Krone, 2003; Mazgajska & Mazgajski, 2020; Sistani et al., 2021), while their primary habitats (steppes and wild river floodplains) have been degraded in many areas, especially in central Europe. It has recently been suggested that green toads tend to continuously moving towards the city centers in contrast to other amphibians, such as the common toad (*Bufo bufo*), which showed the opposite trend (Mazgajska & Mazgajski, 2020). Green toads are known to tolerate higher salt concentrations in their breeding habitats than other amphibians, enabling them to breed in salty steppe lakes and brackish water (Gordon, 1962; Schmidt & Loman, 2019). This feature may also help them to tolerate urban environments that are often polluted with road salt (E. M. Hall et al., 2020). However, a study in Germany indicated that anthropogenic land-use changes are reducing the green toads lifetime (Sinsch et al., 2007), hence, breeding success would need to counteract such effect in order to maintain stable populations.

In Austria the green toad, with only few exceptions, occurs mainly in the East of the country, including remaining steppes and former wild river floodplain areas (Cabela et al., 2001). In our study area in Vienna green toads have been reported from inner districts, as well as the suburbs (Cabela et al., 2003; Sistani et al., 2021; Staufer, 2022; Staufer et al., 2022). However, in-depth analysis of factors contributing to the occurrence of green toads in cities is still lacking, despite of numerous reports on urban populations. The habitat requirements we assume for green toads (preference for open and sunny terrain, and high tolerance for pollution) might be exemplary for a range of species that can seek refuge in city areas. Therefore, by identifying important landscape factors for the green toad, preserving such features will likely increase the overall biodiversity in cities.

To tackle our research question, we used the “Austrian Herpetofauna Database” of the Natural History Museum Vienna and open land-use data from the Viennese local government to model the presence and absence of green toads and thereby reveal land-use dependences of the green toad. While we expected that most urban land-use types have negative effects on green toad presence, we aimed to identify land-use types that positively affect green toad occurrence. This could be used inform urban landscape planning and urban wildlife managing, especially with a focus on endangered animals, that use urban areas as refuges.

## Methods

We used green toad occurrence data (from 2007 to 2020, n = 132 records) obtained from the “Austrian Herpetofauna Database” of the Natural History Museum Vienna. The observations recorded in the database originate from field surveys of the staff of the Herpetological Collection, data transferred by professional herpetologists and Citizen Scientists, scientific publications as well as unpublished investigations, like reports or degree theses, preserved collection specimens, historical indexes, and data exchange with organizations which store regional herpetological inventories. We did not differentiate between different type of reports or number of specimens found; every account of green toads was used as ‘presence’ in our analysis.

All analysis were done in the statistical software R (R Core Team, 2020). To model toad presence, we used a generalized linear model with a binomial error distribution. Following the advice by Barbet-Massin et al. (2012), we generated 10,000 pseudo-absences, which were randomly chosen from the area of Vienna, we adjusted the weights of the data points, in order for the weight sum to be identical between presences (n = 132) and pseudo-absences. Following the approach described in the dismo package (Hijmans et al., 2017) we extracted the sum of the land-use types surrounding each data-point in a radius of 500m, after rasterizing the land-use data (using 25 m^2^ tiles). Land-use data (for 2012) was obtained from the Open Government Data platform (Stadt Wien, 2014). We used the area of each land use type in a 500 m radius of each presence and pseudo-absence as predictors in our model, including all two-way interactions between land-use types. In order to avoid overfitting we performed an automated model selection using the buildmer package (function buildglmmTMB) (Voeten, 2020), using least likelihood ratio tests. The model which performed best was then used to obtain model predictions (using ggpredict from the package ggeffects (Lüdecke, 2018)). Model plots where obtained using the ggeffects plot function with adaptations using ggplot2 (Wickham, 2016) functionality and stitched together using the package patchwork (Pedersen, 2020). The model table was generated using the sjPlot package (Lüdecke, 2020). We plotted the land-use data using the ggplot function. The kernel density map of toad occurrence was created using the function ggmap (Kahle & Wickham, 2013) which uses a 2D kernel density estimation based on the kde2d function in the MASS package (Venables & Ripley, 2002). The background map of Vienna was downloaded using the function get_map and available from OpenStreetMap (OpenStreetMap contributors, 2022).

## Results

Vienna’s land-use type distribution followed the expected pattern with more urban land-use types (e.g., densely populated housing areas) closer to the city core, while agricultural areas and forests were found more frequently farther away from the city center (Fig. 1). Green toads were recorded in most parts of the city except for the northwestern part (Fig. 2). After model selection nine land-use variables remained in the model (Table 1), the three land-use types with the strongest effects were forest, densely populated and transformation/construction site areas (Fig. 2).

**Table 1:** Model results showing the effects of all included factors after model selection contributing to toad occurrence.

| Predictors | Green toad occurrence |  |
| --- | --- | --- |
|  | z | p |
| Forest | -7.25 | <0.001 |
| Dense residential area | -3.80 | <0.001 |
| Transformation area / construction site /material extraction | 3.65 | <0.001 |
| Gardening / orchards | 1.60 | 0.111 |
| Water supply | -0.94 | 0.348 |
| Industry / trade / wholesale including warehouses | -1.39 | 0.166 |
| Sparse residential area | -0.80 | 0.421 |
| Military area | 1.73 | 0.083 |
| Sport and baths / indoor sports facilities | -1.27 | 0.203 |

**Figure 1:**
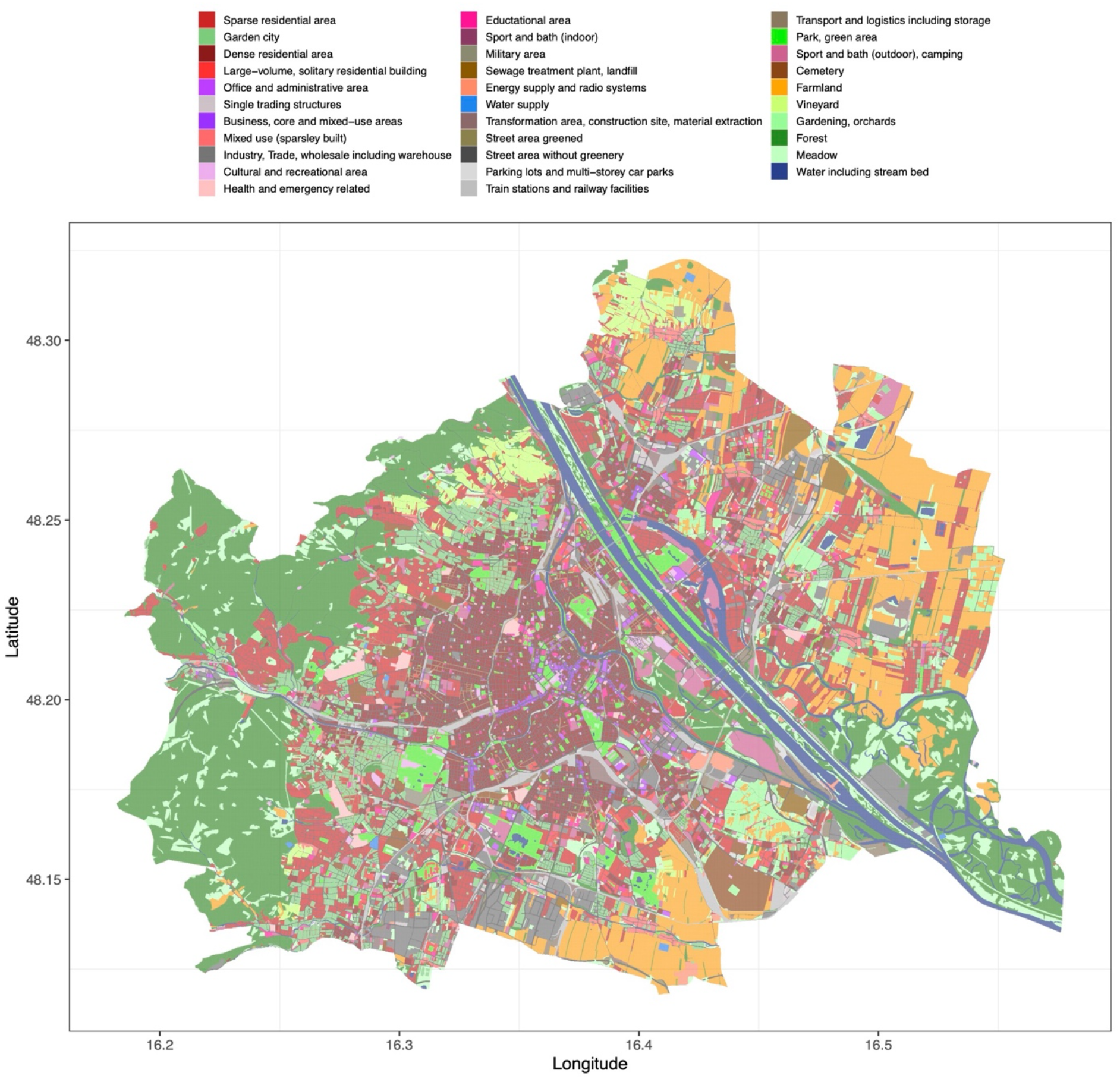
Land use types in Vienna based on the land-use data for 2012 provided by the City of Vienna through the Open Government Data platform (Stadt Wien, 2014).

**Figure 2:**
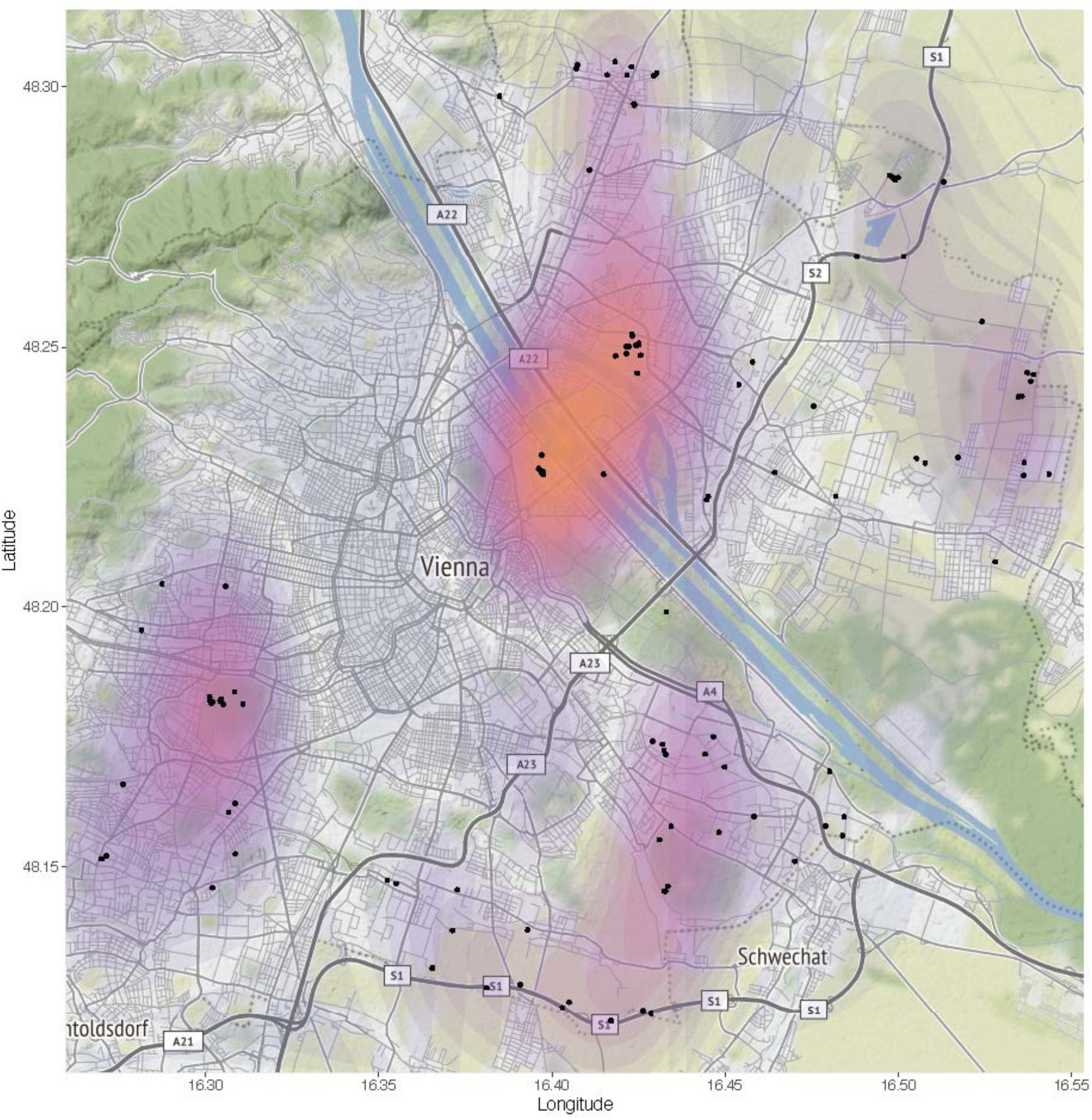
Kernel density representation of green toad occurrence in Vienna. Purple to orange scale corresponding to lower and higher occurrence rate of green toads based on the data base of the Natural History Museum of Vienna (black dots represent toad records).

**Figure 3:**
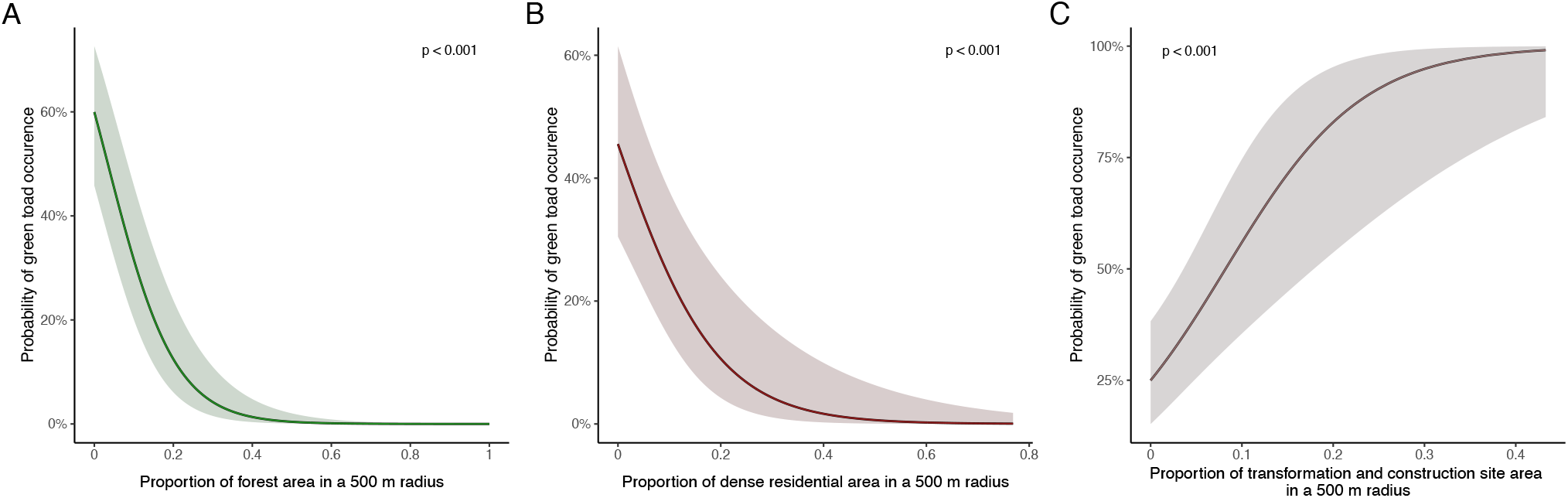
Model predictions of the three significant effects. P-values are shown in the plots. Forest and densely populated areas had strong negative impacts on toad occurrence while transformation/construction sites showed positive effects.

## Discussion

Despite the reported green toad occurrence in Vienna, and other cities, most factors analyzed contributed negatively to the green toad occurrence. Interestingly, the land-use parameter most negatively affecting green toad occurrence was ‘forest’. Forest areas exist in Vienna in several places and of various types, spanning from the woods in the western parts, which are a bit higher in elevation, to the low elevation floodplain related woods around the Danube. Such occurrence pattern is most likely related to the green toads’ preferences of open land (‘steppe species’ (Stöck et al. 2009)), also the more natural floodplain areas where green toads exist (for example around the river Tagliamento), are a highly dynamical environment and have large open sand and gravel areas (Arscott et al., 2002; Kuhn, 2005; Tockner et al., 2003). Such a strong negative impact of a land-use type that is often considered ‘more natural’ or ‘less urban’, is important to consider when planning restoration efforts.

Green toads, as most animal species, were highly significantly negatively impacted by densely populated areas. Densely populated areas can act as a migration/movement barrier and lead to increased mortality (at least) due to roadkill (Glista et al., 2007). Mitigation measures should entail the reduction of traffic in the city, increasing green areas instead of pavement and whenever possible avoiding road construction. In addition, implementation of culverts was shown reduce roadkill incidents of amphibians and other small animals (Dodd et al., 2004; Glista et al., 2009). Given the lack of green toads in densely populated areas, open corridors between existing populations may allow animal movement and reduce the risk of inbreeding (Hinkson & Poo, 2020; Hitchings & Beebee, 1996).

Green toad occurrences were positively affected by the land use category “transformation areas and construction sites”, which is at least partly due to one major long-term construction area in the center of Vienna (“Nordbahnhofgelände”, Stadt Wien 2022a, 2022b), with a well-known green toad population. However, it has been reported from other areas in Europe that green toads can appear quickly at new construction sites, using filled pits as breeding sites (Kühnel & Krone, 2003; Vences et al., 2003). Construction sites, like quarries (Henle et al., 2017; Rogell et al., 2011; Vences et al., 2003), represent open habitats, without dense vegetation or sealed ground. In many ways, it does represent an early succession (i.e., ‘wasteland’) habitat type. However, as technology advances construction sites in cities will be available for increasingly shorter periods. In addition, increasing building activity usually increases ground sealing, and therefore, constitutes a loss of suitable habitats in the long run. Therefore, this would not be a sustainable conservation strategy for green toads (or any other species with such habitat needs). However, preserving such type of habitat, i.e., wastelands or fallows with early succession water bodies, is; although it would involve regular maintenance work in order to prevent further habitat succession (Harper, 2007).

Our analysis, in accordance with published accounts on other species (Darcovich & O’Meara, 2008), shows that species which are not urban exploiters can find a niche in an otherwise dense urban fabric, if certain features are retained. Urban green areas are often dedicated to recuperation for humans and are therefore associated with planting trees, shrubs, flowering plants and in general cool and shaded recreational areas (Hoyle, 2020). However, temporarily unmanaged open land, such as construction sites, fallows and wastelands, can provide essential refuge for endangered species and increase overall biodiversity in cities (Machon, 2021; Mathey & Ring, 2010). The ecological value of such areas might be highly underappreciated, at least in the general public (Hoover et al., 2020). An interesting future avenue of investigation could focus on the community ecology of such ‘wasteland communities’ in cities in response to succession stage, surrounding land use, pollution and global warming (some of the topics already discussed in (Muratet et al., 2007)). Such focus may elucidate the more dynamic aspects of urban wildlife in contrast to the typical urban park and woodland species assemblages. This may influence how policy makers treat and manage open areas (i.e., wastelands) and long-lasting construction sites in cities and provide a way to conserve rare in cities.

## Acknowledgement

We want to thank the numerous people contributing to the database of the herpetological collection of the Natural History Museum of Vienna. Map data copyrighted OpenStreetMap contributors and available from https://www.openstreetmap.org.

## References

Angel, S., Parent, J., Civco, D. L., Blei, A., & Potere, D. (2011). The dimensions of global urban expansion: Estimates and projections for all countries, 2000–2050. Progress in Planning, 75(2), 53–107. https://doi.org/10.1016/j.progress.2011.04.001

Arscott, D. B., Tockner, K., van der Nat, D., & Ward, J. V. (2002). Aquatic Habitat Dynamics along a Braided Alpine River Ecosystem (Tagliamento River, Northeast Italy). Ecosystems, 5(8), 0802–0814. https://doi.org/10.1007/s10021-002-0192-7

Barbet-Massin, M., Jiguet, F., Albert, C. H., & Thuiller, W. (2012). Selecting pseudo-absences for species distribution models: How, where and how many?: How to use pseudo-absences in niche modelling? Methods in Ecology and Evolution, 3(2), 327–338. https://doi.org/10.1111/j.2041-210X.2011.00172.x

Beebee, T. J. C., & Griffiths, R. A. (2005). The amphibian decline crisis: A watershed for conservation biology? Biological Conservation, 125(3), 271–285. https://doi.org/10.1016/j.biocon.2005.04.009

Bogdan, H. (2014). Oradea Zoo – a safe haven for urban Bufotes viridis populations from Oradea (Romania)? Herpetologica Romanica, 8, 33–38.

Bradley, C. A., & Altizer, S. (2007). Urbanization and the ecology of wildlife diseases. Trends in Ecology & Evolution, 22(2), 95–102. https://doi.org/10.1016/j.tree.2006.11.001

Brand, A. B., & Snodgrass, J. W. (2010). Value of artificial habitats for amphibian reproduction in altered landscapes. Conservation Biology, 24(1), 295–301.

Cabela, A., Gressler, S., Teufl, H., & Ellinger, N. (2003). Neu geschaffene Uferstrukturen im Stauraum Freudenau und Folienteiche auf der Wiener Donauinsel: Eine Studie über ihre Wirksamkeit als Trittsteinbiotope für Amphibien. Denisia, 10, 101–142.

Cabela, A., Grillitsch, H., & Tiedemann, F. (2001). Atlas zur Verbreitung und Ökologie der Amphibien und Reptilien in Österreich. Umweltbundesamt, Wien, 1–880.

Collins, J. P., & Storfer, A. (2003). Global amphibian declines: Sorting the hypotheses. Diversity and Distributions, 9(2), 89–98. https://doi.org/10.1046/j.1472-4642.2003.00012.x

Darcovich, K., & O’Meara, J. (2008). An olympic legacy: Green and golden bell frog conservation at Sydney Olympic Park 1993-2006. Australian Zoologist, 34(3), 236– 248. https://doi.org/10.7882/AZ.2008.001

Dodd, C. K., Barichivich, W. J., & Smith, L. L. (2004). Effectiveness of a barrier wall and culverts in reducing wildlife mortality on a heavily traveled highway in Florida. Biological Conservation, 118(5), 619–631. https://doi.org/10.1016/j.biocon.2003.10.011

Gao, J., & O’Neill, B. C. (2020). Mapping global urban land for the 21st century with data-driven simulations and Shared Socioeconomic Pathways. Nature Communications, 11(1), 2302. https://doi.org/10.1038/s41467-020-15788-7

Glista, D. J., Devault, T. L., & Dewoody, J. A. (2007). VERTEBRATE ROAD MORTALITY PREDOMINANTLY IMPACTS AMPHIBIANS. Herpetological Conservation and Biology, 3(1), 77–87.

Glista, D. J., DeVault, T. L., & DeWoody, J. A. (2009). A review of mitigation measures for reducing wildlife mortality on roadways. Landscape and Urban Planning, 91(1), 1–7. https://doi.org/10.1016/j.landurbplan.2008.11.001

Gordon, M. S. (1962). Osmotic regulation in the green toad (Bufo viridis). Journal of Experimental Biology, 39(2), 261–270.

Hall, D. M., Camilo, G. R., Tonietto, R. K., Ollerton, J., Ahrné, K., Arduser, M., Ascher, J. S., Baldock, K. C. R., Fowler, R., Frankie, G., Goulson, D., Gunnarsson, B., Hanley, M. E., Jackson, J. I., Langellotto, G., Lowenstein, D., Minor, E. S., Philpott, S. M., Potts, S. G., … Threlfall, C. G. (2017). The city as a refuge for insect pollinators: Insect Pollinators. Conservation Biology, 31(1), 24–29. https://doi.org/10.1111/cobi.12840

Hall, E. M., Brunner, J. L., Hutzenbiler, B., & Crespi, E. J. (2020). Salinity stress increases the severity of ranavirus epidemics in amphibian populations. Proceedings of the Royal Society B: Biological Sciences, 287(1926), 20200062. https://doi.org/10.1098/rspb.2020.0062

Hamer, A. J., & McDonnell, M. J. (2008). Amphibian ecology and conservation in the urbanising world: A review. Biological Conservation, 141(10), 2432–2449. https://doi.org/10.1016/j.biocon.2008.07.020

Harper, C. A. (2007). Strategies for Managing Early Succession Habitat for Wildlife. Weed Technology, 21(4), 932–937.

Hassell, J. M., Begon, M., Ward, M. J., & Fèvre, E. M. (2017). Urbanization and Disease Emergence: Dynamics at the Wildlife–Livestock–Human Interface. Trends in Ecology & Evolution, 32(1), 55–67. https://doi.org/10.1016/j.tree.2016.09.012

Henle, K., Dubois, A., & Vershinin, V. (2017). Mass anomalies in green toads (<i>Bufotes viridis>/i>) at a quarry in Roßwag, Germany: Inbred hybrids, radioactivity or an unresolved case? Mertensiella, 25, 185–242.

Hijmans, R. J., Phillips, S., Leathwick, J., & Elith, J. (2017). dismo: Species Distribution Modeling (R package version 1.1-4) [Computer software]. https://CRAN.R-project.org/package=dismo

Hinkson, K. M., & Poo, S. (2020). Inbreeding depression in sperm quality in a critically endangered amphibian. Zoo Biology, 39(3), 197–204. https://doi.org/10.1002/zoo.21538

Hitchings, S. P., & Beebee, T. J. C. (1996). Genetic substructuring as a result of barriers to gene flow in urban Rana temporaria (common frog) populations: Implications for biodiversity conservation. Heredity, 79, 117–127.

Hoover, D. L., Bestelmeyer, B., Grimm, N. B., Huxman, T. E., Reed, S. C., Sala, O., Seastedt, T. R., Wilmer, H., & Ferrenberg, S. (2020). Traversing the Wasteland: A Framework for Assessing Ecological Threats to Drylands. BioScience, 70(1), 35–47. https://doi.org/10.1093/biosci/biz126

Hoyle, H. (2020). What Is Urban Nature and How Do We Perceive It? In Naturally Challenged: Contested Perceptions and Practices in Urban Green Spaces (S. 9–36). Springer.

IUCN. (2021). The IUCN Red List of Threatened Species. IUCN Red List of Threatened Species. https://www.iucnredlist.org/en

Kahle, D., & Wickham, H. (2013). ggmap: Spatial Visualization with ggplot2. 5, 18.

Konowalik, A., Najbar, A., Konowalik, K., Dylewski, Ł., Frydlewicz, M., Kisiel, P., Starzecka, A., Zaleśna, A., & Kolenda, K. (2020). Amphibians in an urban environment: A case study from a central European city (Wrocław, Poland). Urban Ecosystems, 23(2), 235–243. https://doi.org/10.1007/s11252-019-00912-3

Kuhn, K. (2005). Die Kiesbänke des Tagliamento (Friaul, Italien)-Ein Lebensraum für Spezialisten im Tierreich. Jahrbuch des Vereins zum Schutz der Bergwelt, 70, 37–44.

Kühnel, K.-D., & Krone, A. (2003). Bestandssituation, Habitatwahl und Schutz der Wechselkröte (Bufo viridis) in Berlin – Grundlagenuntersuchungen für ein Artenhilfsprogramm in der Grossstadt. Mertensiella, 14, 299–315.

Luck, G. E., & Smallbone, L. T. (2010). Species diversity and urbanization: Patterns, drivers and implications. In K. J. Gaston (Hrsg.), Urban Ecology (S. 88–119). Cambridge University Press.

Lüdecke, D. (2018). ggeffects: Tidy data frames of marginal effects from regression models. Journal of Open Source Software, 3(26), 772. https://doi.org/10.21105/joss.00772

Lüdecke, D. (2020). sjPlot: Data visualization for statistics in social science (R package version 2.8.3) [Computer software].

Machon, N. (2021). Urban Wastelands Can Be Amazing Reservoirs of Biodiversity for Cities. In F. Di Pietro & A. Robert (Hrsg.), Urban Wastelands (S. 3–18). Springer International Publishing. https://doi.org/10.1007/978-3-030-74882-1_12

Marzluff, J. M. (2001). Worldwide urbanization and its effects on birds. In J. M. Marzluff, R. Bowman, & R. Donnelly (Hrsg.), Avian Ecology and Conservation in an Urbanizing World (S. 19–47). Springer US. https://doi.org/10.1007/978-1-4615-1531-9_2

Mathey, J., & Ring, D. (2010). Urban Wastelands – A Chance for Biodiversity in Cities? Ecological Aspects, Social Perceptions and Acceptance of Wilderness by Residents. In N. Müller, P. Werner, & J. G. Kelcey (Hrsg.), Urban Biodiversity and Design (Conservation Science and Practice) (S. 406–424). Wiley-Blackwell.

Mazgajska, J., & Mazgajski, T. D. (2020). Two amphibian species in the urban environment: Changes in the occurrence, spawning phenology and adult condition of common and green toads. The European Zoological Journal, 87(1), 170–179. https://doi.org/10.1080/24750263.2020.1744743

McKinney, M. L. (2002). Urbanization, Biodiversity, and Conservation: The impacts of urbanization on native species are poorly studied, but educating a highly urbanized human population about these impacts can greatly improve species conservation in all ecosystems. BioScience, 52(10), 883–890. https://doi.org/10.1641/0006-3568(2002)052[0883:UBAC]2.0.CO;2

McKinney, M. L. (2008). Effects of urbanization on species richness: A review of plants and animals. Urban Ecosystems, 11(2), 161–176. https://doi.org/10.1007/s11252-007-0045-4

Moore, M., Gould, P., & Keary, B. S. (2003). Global urbanization and impact on health. International Journal of Hygiene and Environmental Health, 206(4–5), 269–278. https://doi.org/10.1078/1438-4639-00223

Muratet, A., Machon, N., Jiguet, F., Moret, J., & Porcher, E. (2007). The Role of Urban Structures in the Distribution of Wasteland Flora in the Greater Paris Area, France. Ecosystems, 10(4), 661. https://doi.org/10.1007/s10021-007-9047-6

Murray, M. H., Sánchez, C. A., Becker, D. J., Byers, K. A., Worsley-Tonks, K. E., & Craft, M. E. (2019). City sicker? A meta-analysis of wildlife health and urbanization. Frontiers in Ecology and the Environment, 17(10), 575–583. https://doi.org/10.1002/fee.2126

Ofori, B. Y., Garshong, R. A., Gbogbo, F., Owusu, E. H., & Attuquayefio, D. K. (2018). Urban green area provides refuge for native small mammal biodiversity in a rapidly expanding city in Ghana. Environmental Monitoring and Assessment, 190(8), 480. https://doi.org/10.1007/s10661-018-6858-1

OpenStreetMap contributors. (2022). Planet dump retrieved from https://planet.osm.org. https://www.openstreetmap.org

Pedersen, T. L. (2020). patchwork: The composer of plots (R package version 1.1.1) [Computer software].

R Core Team. (2020). R: A language and environment for statistical computing. R Foundation for Statistical Computing. https://www.R-project.org/

Rannap, R., Lõhmus, A., & Jakobson, K. (2007). Consequences of coastal meadow degradation: The case of the natterjack toad (Bufo Calamita) in Estonia. Wetlands, 27(2), 390. https://doi.org/10.1672/0277-5212(2007)27[390:COCMDT]2.0.CO;2

Rogell, B., Berglund, A., Laurila, A., & Höglund, J. (2011). Population divergence of life history traits in the endangered green toad: Implications for a support release programme. Journal of Zoology, 285(1), 46–55. https://doi.org/10.1111/j.1469-7998.2011.00843.x

Scheffers, B. R., & Paszkowski, C. A. (2011). The effects of urbanization on North American amphibian species: Identifying new directions for urban conservation. Urban Ecosystems, 15(1), 133–147.

Schmidt, A., & Loman, J. (2019). Salt tolerance of Bufotes viridis eggs and tadpoles. Alytes, 37(1–2), 46–62.

Sinsch, U., Leskovar, C., Drobig, A., König, A., & Grosse, W.-R. (2007). Life-history traits in green toad (Bufo viridis) populations: Indicators of habitat quality. Canadian Journal of Zoology, 85(5), 665–673.

Sistani, A., Burgstaller, S., Gollmann, G., & Landler, L. (2021). The European green toad, Bufotes viridis, in Donaufeld (Vienna, Austria): Status and size of the population. Herpetozoa, 34, 259–264. https://doi.org/10.3897/herpetozoa.34.e75578

Soanes, K., & Lentini, P. E. (2019). When cities are the last chance for saving species. Frontiers in Ecology and the Environment, 17(4), 225–231. https://doi.org/10.1002/fee.2032

Stadt Wien. (2014). General data on actual land use in Vienna, based on an aerial interpretation and supplementary facts. [Land use data]. https://data.wien.gv.at/daten/geo?service=WFS&request=GetFeature&version=1.1.0&typeName=ogdwien:REALNUT2012OGD&srsName=EPSG:4326&outputFormat=shape-zip

Stadt Wien. (2022a). Rudolf-Bednar-Park. https://www.wien.gv.at/english/environment/parks/bednar.html

Stadt Wien. (2022b). Stadtentwicklungsgebiet Nordbahnhof—Projektübersicht. https://www.wien.gv.at/stadtentwicklung/projekte/nordbahnhof/projekte/index.html

Staufer, M. (2022). Die Wechselkröten der Simmeringer Haide in Wien. ÖGH-Aktuell, 60, 29– 35.

Staufer, M., Burgstaller, S., & Landler, L. (2022). Beitrag zur Phänologie der Wechselkröte in Wien: Laichbeginn in den Jahren 2019 und 2020. ÖGH-Aktuell, 60, 36–37.

Suislepp, K., Rannap, R., & Lõhmus, A. (2011). Impacts of artificial drainage on amphibian breeding sites in hemiboreal forests. Forest Ecology and Management, 262(6), 1078– 1083. https://doi.org/10.1016/j.foreco.2011.06.001

Tockner, K., Ward, J. V., Arscott, D. B., Edwards, P. J., Kollmann, J., Gurnell, A. M., Petts, G. E., & Maiolini, B. (2003). The Tagliamento River: A model ecosystem of European importance. Aquatic Sciences - Research Across Boundaries, 65(3), 239–253. https://doi.org/10.1007/s00027-003-0699-9

Van Helden, B. E., Close, P. G., Stewart, B. A., Speldewinde, P. C., & Comer, S. J. (2021). Critically Endangered marsupial calls residential gardens home. Animal Conservation, 24(3), 445–456. https://doi.org/10.1111/acv.12649

Venables, W. N., & Ripley, B. D. (2002). Modern Applied Statistics with S Fourth edition by, World.

Vences, M., Glaw, F., & Franzen, M. (2003). Perspektiven für den kostengünstigen Erhalt von Lebensräumen in Abgrabungen und ihre Bedeutung für die Wechselkröte (Bufo viridis). Mertensiella, 14, 316–327.

Vlahov, D. (2002). Urbanization, Urbanicity, and Health. Journal of Urban Health: Bulletin of the New York Academy of Medicine, 79(90001), 1S – 12. https://doi.org/10.1093/jurban/79.suppl_1.S1

Voeten, C. C. (2020). buildmer: Stepwise Elimination and Term Reordering for Mixed-Effects Regression (R package version 1.5.) [Computer software]. https://CRAN.R-project.org/package=buildmer

Wickham, H. (2016). ggplot2: Elegant graphics for data analysis. Springer.

